# A conformational change in the N terminus of SLC38A9 signals mTORC1 activation

**DOI:** 10.1101/339937

**Authors:** Ma Jinming, Hsiang-Ting Lei, Tamir Gonen

## Abstract

mTORC1 is a central signal hub that integrates multiple environmental cues, such as cellular stresses, energy levels, nutrients and certain amino acids, to modulate metabolic status and cellular responses. Recently, SLC38A9, a lysosomal amino acid transporter, has emerged as a sensor for luminal arginine levels and as an activator of mTOCRC1. The activation of mTORC1 occurs through the N-terminal domain of SLC38A9. Here, we determined the crystal structure of SLC38A9 and surprisingly found its N-terminal fragment inserted deep into the transporter, bound in the substrate binding pocket where normally arginine would bind. Compared with our recent arginine bound structure of SLC38A9, a significant conformational change of the N-terminal domain was observed. A ball-and-chain model is proposed for mTORC1 activation where in the starved state the N-terminal domain of SLC38A9 is buried deep in the transporter but in the fed state the N-terminal domain could be released becoming free to bind the Rag GTPase complex and to activate mTORC1. This work provides important new insights into how SLC38A9 senses the fed state and activates the mTORC1 pathways in response to dietary amino acids.

**One Sentence Summary:** N-plug inserted state of SLC38A9 reveals mechanisms of mTORC1 activation and arginine-enhanced luminal amino acids efflux.

## Main Text

The mechanistic target of rapamycin complex 1 (mTORC1) protein kinase acts as a central signaling hub to control cell growth and balance the products from anabolism and catabolism *(1–3)*. Not surprisingly this pathway is dysregulated in many diseases *(4, 5)*. Activation of the mTORC1 is mediated by a variety of environmental cues such as nutrients, cellular stresses and energy levels *(6, 7)*. Specifically, certain amino acids signal to mTORC1 through two Ras-related guanosine triphosphatases (GTPases) *(8, 9)*. When amino acids are abundant, the heterodimeric Rag GTPases adopt an active state and promote the recruitment of mTORC1 to the lysosomal surface *(10)*, which is now recognized as a key subcellular organelle involved in mTORC1 regulation *(11)*. Several essential amino acids in the lysosomal lumen including arginine, leucine and glutamine have been identified as effective activators of mTORC1 *(12–15)*. However, the molecular basis of the amino acids sensing mechanism has remained, by and large, elusive. Recently, SLC38A9, a low-affinity arginine transporter on lysosome vesicles, was identified as a direct sensor of lumen arginine levels for the mTORC1 pathway *(16–18)*. SLC38A9 also mediates the efflux of essential amino acids from lysosomes, such as leucine, in an arginine regulated manner *(19)*, to drive cell growth by modulating cytosolic sensors *(20, 21)*. Moreover, SLC38A9 senses the presence of luminal cholesterol and activates mTORC1 independently of its arginine transport *(22)*.

SLC38A9 is a transceptor. Studies showed that two parts of SLC38A9, its N-terminal domain and its transmembrane bundle, are responsible for two distinct functions. The bulk of SLC38A9 are ll alpha helices that pack against one another forming a transmembrane bundle that transports amino acids and function as an amino acid transporter *(23)*. The N-terminus of SLC38A9, om the other hand, was previously shown to interact directly with the Rag-Regulator complex to activate mTORC1 *(16)*. Collectively, these results suggest that SLC38A9 is a “transceptor”, which is membrane protein that embodies the functions of both a transporter and a receptor *(23–27)*.

Recently we solved the crystal structure of N-terminally truncated SLC38A9 from *Danio rerio* (ΔN-drSLC38A9) with arginine bound *(23)*. The substrate arginine was observed deep in the transporter at a binding pocket consisting of residues from TM1a, TM3 and TM8 of SLC38A9. Because the N-terminally truncated form of SLC38A9 was used that initial study focused solely on the transporter function of SLC38A9 and resulting structures could not inform on the signaling function of SLC38A9.

Here we report a new crystal structure of drSLC38A9 with its N-terminus but without the substrate arginine. Surprisingly, we found that part of the N-terminus formed a beta hairpin that lodged itself deep in the transporter occupying the arginine binding site and blocking the transport path. These new results suggest that in the fed state the N-terminal domain would be released from within SLC38A9 and freed to interact with the Rag GTPase and activate mTORC1. We propose a ball-and-chain model to describe this mechanism of amino acid sensation and signaling by SLC38A9.

In the present study we used the antibody fragment llD3 to facilitate crystallization of SLC38A9 in the absence of substrate. Well-ordered crystals were diffracted to ~3.4 Å with high completeness and acceptable refinement statistics (Table. S1). Each asymmetric unit contained two copies of drSLC38A9-Fab complex, arranged in a propeller-like head-to-head fashion (fig. S1). As with the recently determined structure *(23)*, the transmembrane domain of drSLC38A9 was captured in the cytosol-open state and was folded into the same inverted topology repeats made of TMs 1-5 and TMs 6-10 with TMll wrapping around the transceptor (Fig. 1A and 1B). The two structures shared an overall similar fold with an r.m.s.d. of 0.8 Å. However, instead of an arginine molecule bound, this time an unexpected electron density was observed, which extended along the solvent accessible tunnel leading from the substrate binding site to the cytosolic side of SLC38A9 (fig. S2). The density was of sufficient quality to allow an unambiguous assignment of drSLC38A9 N-terminal section from Asp 75 to Leu 9l (fig. S3). This fragment formed a folded domain, resembling a beta hairpin, filling the entire path from the cytosolic side of SLC38A9 to the substrate binding site (Fig. 1C). Electrostatic potential analysis indicated that the transport pathway in SLC38A9 is generally positively charged, while the N-terminal fragment (referred to as the “N-plug” from this point on) is largely negatively charged (fig. S4), suggesting that the interaction is electrostatically driven. Deletion of the N-terminal domain of drSLC38A9 did not affect arginine transport (Fig. 1D), indicating that the N-plug does not directly participate in arginine translocation.

**Fig. 1.**
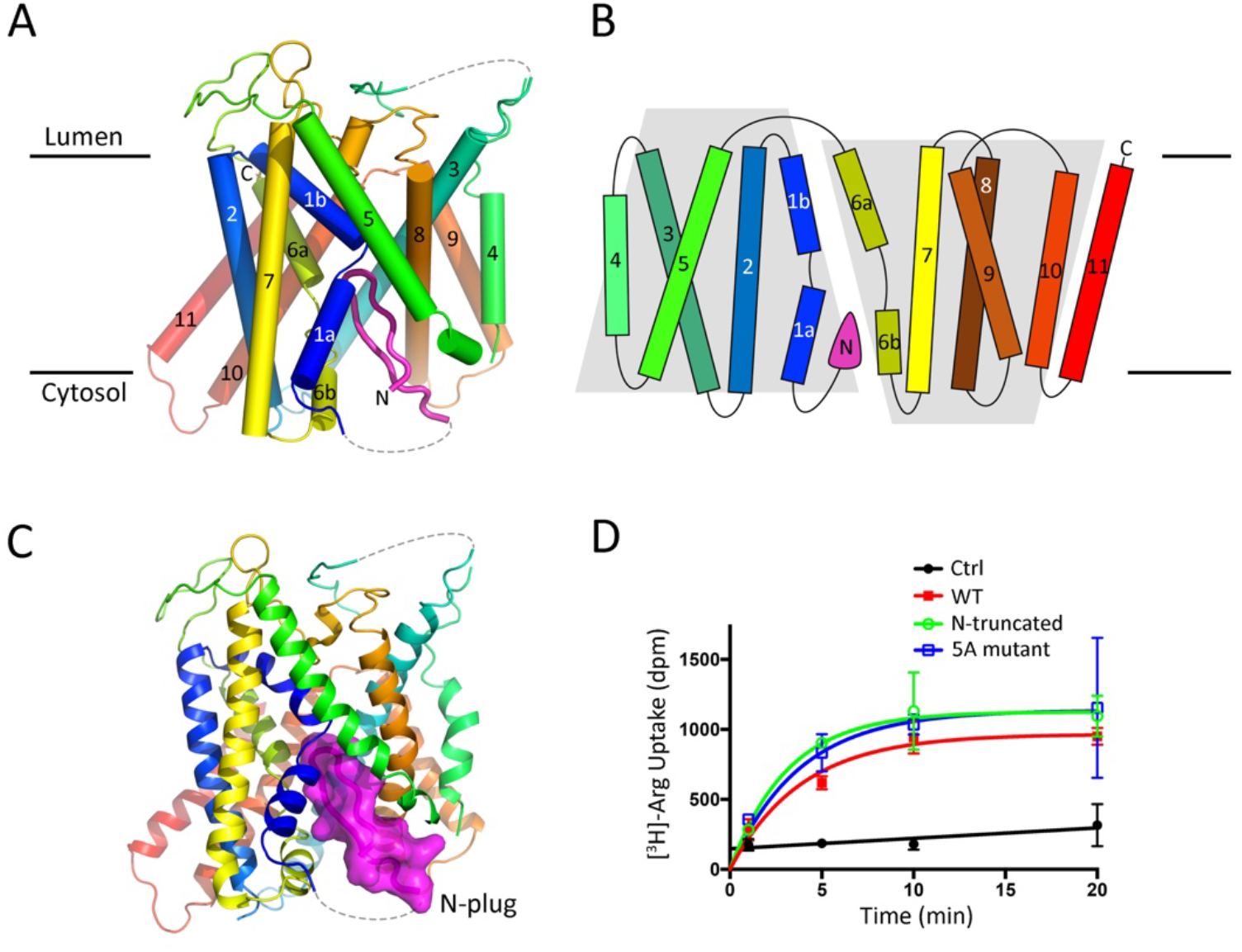
Structure of drSLC38A9 in the N-plug inserted state. (**A**) Stereo view in the plane of the membrane. TMs are rainbow colored as blue to red from N- to C-terminus. The N-plug is shown in magenta. (**B**) Two-dimensional topology model of drSLC38A9, which is folded into a characteristic 2-fold LeuT-like pseudo-symmetry (five transmembrane-helix inverted-topology repeat). N-plug is marked by a filled pink triangle, next to the TM1a helix. (**C**) The N-plug blocks an otherwise cytosol-open state of drSLC38A9. (**D**) Truncation or mutation of the N-plug does not affect arginine transport. Shown here is the time course of [^3^H]-arginine uptake in proteoliposomes reconstituted with purified wild-type drSLC38A9 and its mutants. Error bars represent standard error of the mean (s.e.m.) of triplicate experiments.

We captured SLC38A9 in a new state that we term the “N-plug inserted state”. TMs 1, 5, 6 and 8 of SLC38A9 form a V-shaped cavity into which the N-plug inserts and is stabilized by several bonds (Fig. 2). At the tapered tip on the N-plug, Ser 80 and His 81 bound to the main-chain carbonyl oxygens of Thr 117, Met 118 and Met 119 in the unwound region of TM1 (Fig. 2A). His 81 further stabilizes the tip region of the N-plug through a hydrogen bond between its imidazole side chain and Thr 121 (Fig. 2A). Likewise, the main-chain carbonyl oxygen of Ile 84 is bound to Cys 363 on TM6 (Fig. 2B). At this juncture, the N-plug is jammed in between the two essential TMs 1 and 6 where it would probably prevent the transmembrane domain from transitioning to an alternate state for transport. At the N-terminus of the N-plug, the flanking residues are anchored against TM5 through a hydrogen bond formed between the main-chain carbonyl oxygens of Val 77 and the side-chain hydroxyl group of Thr 303 (Fig. 2C). At the C-terminus of the N-plug, the Tyr-Ser pairs involving Tyr 87, Tyr 448 and Ser 88, Ser 297 also stabilize the interaction by hydrogen bonds (Fig. 2D). All residues that participate in the inter-domain interactions are conserved across species as indicated in the sequence alignments (fig. S3), suggesting that this interaction is evolutionarily conserved and likely plays an important functional role. The beta-hairpin structure of the N-plug is also self-stabilized by several hydrogen bonds between Ser 80 and Glu 82, His76 and Tyr 85 which fasten the two ends of the N-plug together (Fig. 2E). Structural modeling by PEP-FOLD *(28, 29)* indicated that the beta hairpin motif would be converted to an alpha helical fragment should these residues were changed to alanines (fig. S5).

**Fig. 2.**
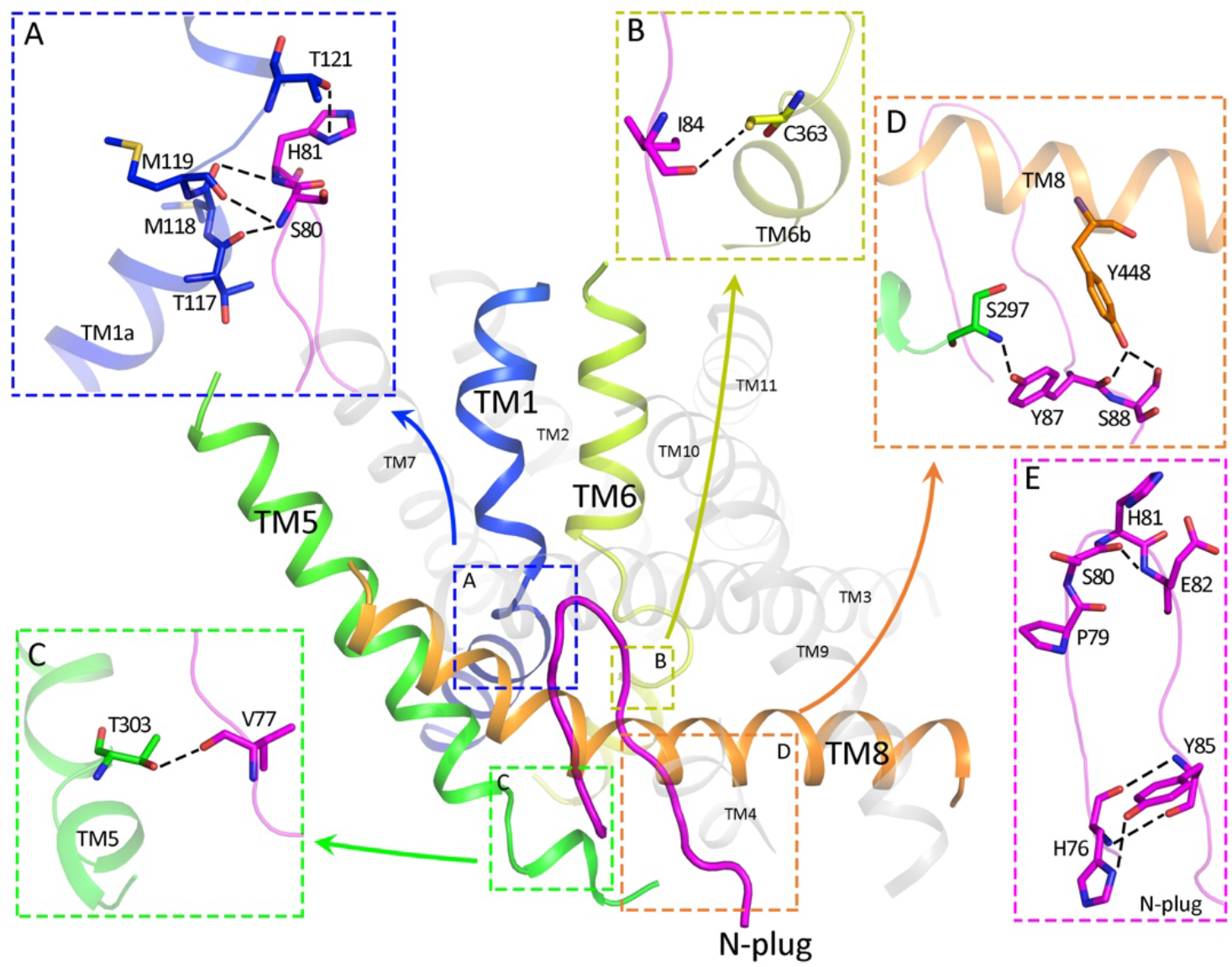
Inter-and intra-domain interactions of the N-plug inside SLC38A9. (**A** to **D**) The N-plug interacts with transmembrane bundle though multiple inter-domain hydrogen bonds. Residues that contribute interactions between the N-plug and TMs are highlighted in sticks, of which hydrogen bonds are depicted as dashed lines. (**E**) The folded conformation of N-plug as a beta hairpin is complementarily stabilized by several intra-domain interactions.

SLC38A9 has higher affinity toward leucine than arginine although the transport of leucine is largely facilitated by the presence of arginine *(19)*. Uptake studies performed here with drSLC38A9 corroborate the previous findings using the human protein (Fig. 3B). Leucine uptake was significantly higher in the presence of supplemented arginine than without. An overlay of the N-plug bound structure and the arginine bound structure indicated that the same set of backbone atoms are used for binding the N-plug and the arginine molecule (Fig. 3A). This superposition suggests that in the presence of arginine the N-terminal plug may not occupy the binding site, but that in the absence of arginine it would be free to insert and bind. Is it possible, therefore, that in the presence of arginine the released N-terminal plug could play an important role in facilitating leucine transport?

**Fig. 3.**
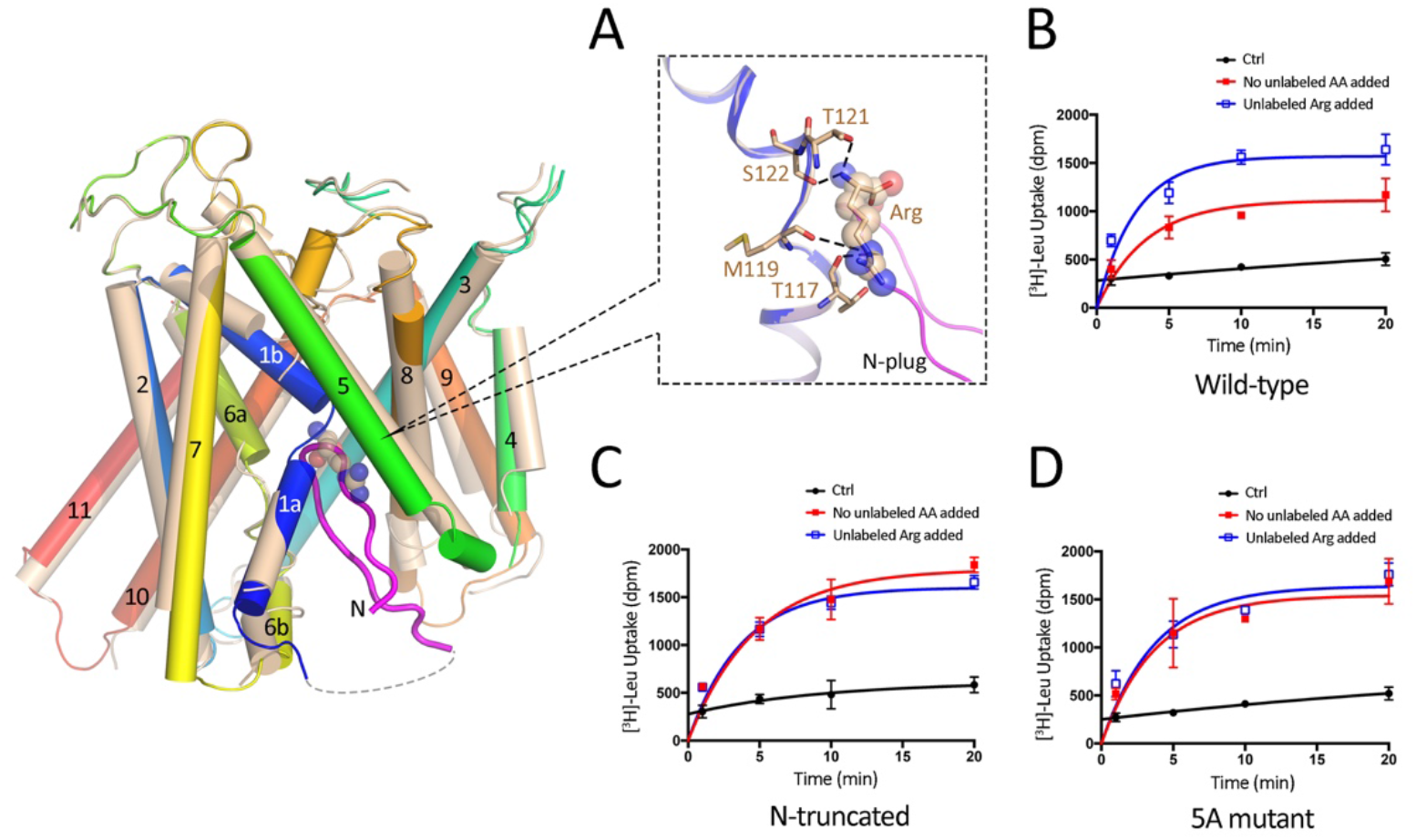
The N-plug is essential for arginine enhanced transport of leucine by drSLC38A9. (**A**) Superposition of substrate binding site of arginine-bound state (PDB ID: 6C08) with N-plug inserted state of drSLC38A9. TM1 of two different states are shown in gold and blue, respectively. Atoms of arginine molecule are depicted as spheres while the N-plug in magenta. (**B**) Adding 200μm unlabeled arginine boosts leucine transport by wild-type drSLC38A9 in proteoliposomes. (**C** and **D**) Either deletion or mutation of N-plug interferes the arginine enhancement of leucine transport. Without adding supplemented arginine, the mutant proteins show similar transport capacity for leucine regardless whether arginine was supplemented.

To examine whether the N-terminal plug plays an important role in facilitating leucine transport, two drSLC38A9 variants were generated. One has residues 1-96 of N-terminus deleted (called N-truncated). The other has 5 key residues mutated (P79A, S80A, H81A, E82A, and Y85A) in the N-plug (named 5A mutant) which would lead to a disrupted secondary structure of the N-plug (fig. S5). As observed in the uptake study, both variants could transport arginine like the wild-type drSLC38A9 even with the dramatic structural changes at the N-plug (Fig. 1D). From the results of leucine uptake by wild-type drSLC38A9, the arginine-enhanced transport of leucine is reflected as increased uptake of [^3^H]-leucine when the buffer was supplemented with arginine. This characteristic of arginine-enhanced leucine transport was lost when the N-plug was eliminated or its structure altered by mutation (Fig. 3C and 3D). Only the SLC38A9 with intact N-terminal plug in its native beta hairpin like structure showed the characteristic enhanced leucine uptake in the presence of supplemented arginine (Fig. 3B).

It is known that the N-terminal domain of SLC38A9 can bind to, and activate, the Rag GTPases complex *(16)*. Moreover, it was shown that the N-terminal fragment of human SLC38A9 (hSLC38A9) was sufficient and required to bind the Ragulator-Rag GTPases complex *(16)*. The binding of Rag GTPases and the human SLC38A9 involves the 85PDH87 motif *(17)*, Pro 85 and Pro 90 *(16)*, corresponding to a conserved region on the N-plug in drSLC38A9 (fig. S3). To probe the N-plug interaction with the Rag GTPases in drSLC38A9, we co-purified zebrafish Rag GTPases complex (drRagA and drRagC) with two N-terminal fragments of drSLC38A9 by size-exclusion chromatography. The first fragment (residues 1-96) contained the N terminus in its entirety (called drSLC38A9-N.1) while in the second fragment (residues 1-70) the N-plug was deleted (called drSLC38A9-N.2). Fractions from size exclusion chromatography were collected and analyzed by SDS-PAGE (fig. S6). Contrary to fragment drSLC38A9-N.1 which maintains the N-terminal domain in its entirety, the N-plug deleted construct, drSLC38A9-N.2 did not associate with Rag GTPases complex (fig. S6). These results clearly demonstrated that the interaction between the zebrafish SLC38A9 N-terminus and the zebrafish Rag-GTPase recapitulate the experiments reported previously using the human proteins *(17, 16)*: the same region of the N-plug of drSLC38A9 is essential for binding with Rag GTPases complex.

In considering our recently determined structure of SLC38A9 with arginine bound, and the current structure without arginine but with the N-plug inserted into the arginine binding site, we now captured SLC38A9 is at least two distinct conformations of the N-terminus. The first is when the N-plug is bound snuggly in the arginine binding site (in the absence of arginine, starved state) and the second where the N-terminal plug was released and the substrate binding site was occupied by arginine (in the presence of arginine, fed state). The vestibule into which the N-terminal plug inserts measures ~20Å in diameter. A recently determined crystal structure of Rag GTPases-Ragulator *(30–32)* indicated that the GTPases-regulator is far too large to fit inside the vestibule of SLC38A9 suggesting that the N-plug must exit the tranceptor for binding the Rag GTPase. Together, these data suggest a mechanism by which SLC38A9 can act as a receptor to signal the activation of Rag GTPase and therefore of mTORC1 in the presence of arginine.

We thus propose a ball-and-chain model (Fig. 4). At lower arginine concentrations, two conformational states could be at an equilibrium where the N-terminal plug is equally inserted or released from the arginine binding site of SLC38A9. When the equilibrium shifts to the right in the fed state with elevated arginine levels, an arginine molecule will occupy the binding site of SLC38A9 for transport and the N-terminal plug would remain released as long as arginine flows. As a result, the N-terminal plug becomes available for binding to the Rag GTPases complex which in turn could activate the mTORC1. Moreover, the release of the N-terminal plug from the helical bundle of SLC38A9 will also facilitate the efflux of other essential amino acids, which simultaneously increases the cytosolic concentration of amino acids and synergistically activates mTORC1 through other cytosolic sensors.

**Fig. 4.**
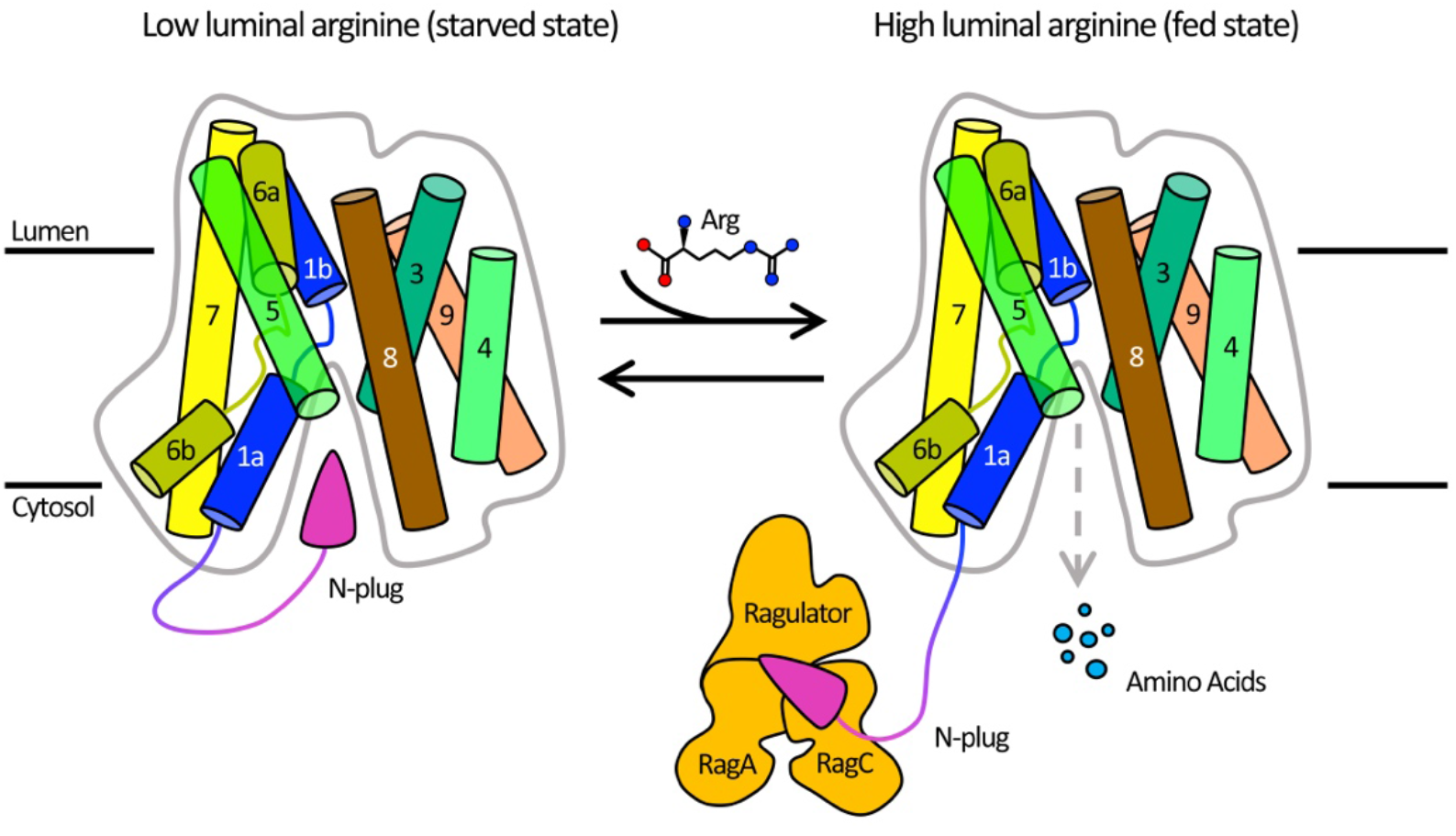
Ball and chain model of SLC38A9 for mTORC1 activation and amino acid transport. At low luminal arginine, N-plug domain naturally samples both the inserted and released state as an equilibrium. As the concentration of luminal arginine increase in the fed state, arginine molecules enter the substrate binding site and the N-plug remains in the released state while arginine transport takes place. In the released state the N-plug could both trigger the efflux of other luminal amino acids such as leucine and interact with the Rag-GTPases to activate the mTORC1 signaling pathway.

While the present study provides the first line of evidence on the function of SLC38A9 as a transporter and sensor for amino acids it remains unclear how the N-terminal domain associates with the Rag GTPase complex. Likewise, it is still not known what the open-to-lumen conformation of the transporter looks like and whether the N-plug remains inserted or not. Future studies must delve into these important open questions but with the newly proposed ball-and-chain model for signaling new biochemical assays can be designed and tested.

## Acknowledgements

We thank D. Cawley for development and production of monoclonal antibodies; K. Rajashankar and the staffs in Northeastern Collaborative Access Team (NE-CAT) for their support with X-ray data collection; J. Hattne for discussions over structural determination; L. Shao for careful review and scientific feedback on the manuscript. This work is based upon research conducted at the NE-CAT beamlines, which are funded by the National Institute of General Medical Sciences from the National Institutes of Health (P41 GM103403). The Pilatus 6M detector on 24-ID-C beam line is funded by a NIH-ORIP HEI grant (S10 RR029205). This research used resources of the Advanced Photon Source, a U.S. Department of Energy (DOE) Office of Science User Facility operated for the DOE Office of Science by Argonne National Laboratory under Contract No. DE-AC02-06CHll357. Research in the Gonen laboratory is funded by the Howard Hughes Medical Institute. The coordinates and the structure factors have been deposited in the Protein Data Bank (PDB) under accession codes 6DCI.

## Supplementary Materia

### Materials and Methods

#### Protein expression and purification

The gene of wild-type SLC38A9 (NP_001073468.1) from *Danio rerio* and its site-directed mutants were produced by polymerase chain reaction (PCR) and then subcloned into a pFastbacl vector containing an octa-histidine tag with a thrombin-cleavage site at the N-terminus. drSLC38A9 protein and its variants were overexpressed in *Spodoptera frugiperda* Sf-9 insect cells following the protocol of Bac-to-Bac Baculovirus Expression System (Invitrogen). Cells were harvested at 60 hours after infection and homogenized in the low salt buffer containing 20 mM Tris pH 8.0, 150 mM NaCl supplemented with cOmplete Protease Inhibitor Cocktail (Roche). The lysate was collected and ultra-centrifuged at 130,000 *×g* for 1 hour. Pelleted membrane was then resuspended and washed with the high salt buffer containing 1.0 M NaCl and 20 mM Tris (8.0) followed by ultracentrifugation. The pellets were resuspended in the low salt buffer, frozen in liquid nitrogen and stored in −80°C until further use.

To purify drSLC38A9 protein and its variants, membrane fraction was thawed and solubilized with 2% n-dodecyl-b-D-maltopyranoside (DDM, Anatrace) in 20 mM Tris pH 8.0, 500 mM NaCl, 5% glycerol, and 0.2% Cholesteryl Hemisuccinate Tris Salt (CHS, Anatrace) for 4 hours at 4 °C. Following another ultra-centrifugation at 130,000 *×g* for 1 hour, the supernatant was loaded onto TALON Metal Affinity Resin (Clontech) and incubated at 4°C overnight. The resins were washed by 5× column volumes of 50 mM imidazole, 20 mM Tris pH 8, 500 mM NaCl, 0.1% DDM before equilibration in 20 mM Tris pH 8.0, 500 mM NaCl, 0.4% decyl-b-D-maltoside (DM) and 0.02% DDM. The N-terminal octa-histidine tag was removed by in-column thrombin digestion overnight at enzyme:protein molar ratio of 1:1000. The cleaved drSLC38A9 proteins collected in flow-through were then flash frozen in liquid nitrogen and stored in −80°C until use.

#### Fab fragments production

Fab fragments were produced at Monoclonal Antibody Core of Vaccine and Gene Therapy Institute, OHSU. Mouse IgG monoclonal antibodies against drSLC38A9 were raised by standard protocol *(33)* using purified protein in the buffer containing 20 mM Tris pH 8.0, 150 mM NaCl, 0.02% DDM, 0.002% CHS as antigen. Western blot and native-to-denature ELISA assays *(34)* were performed to assess the binding affinity and specificity of the antibodies generated from hybridoma cell lines. Several monoclonal antibodies showing high binding affinity and specificity to conformational epitope were then selected and purified from the hybridoma supernatants. Fab fragments were generated by Papain (Thermo Fisher Scientific) digestion and purified by Protein A affinity chromatography (GE Healthcare) in 20 mM Sodium phosphates pH 8.0, 150 mM NaCl.

#### Purification of drSLC38A9-Fab complexes for crystallization

Purified drSLC38A9 proteins was mixed with excess Fab fragments at a molar ratio of 1:2 for 2 hour, and the mixture was subjected to gel filtration (Superdex 200 Increase 10/300 GL, GE Healthcare) in the buffer containing 20 mM Tris-HCl pH 8.0, 150 mM NaCl and 0.2% DM. The peak fractions containing appropriate drSLC38A9-Fab complexes were then pooled and concentrated to 5 mg/mL for crystallization.

#### Crystallization

Crystallization was carried out by hanging-drop vapor diffusion at 4 °C. Initial hits of drSLC38A9 were identified in multiple conditions containing PEG 400. However, these crystals gave anisotropic diffraction to ~6 Å. Well diffracting crystals were only obtained when drSLC38A9 was co-crystallized as a complex with Fab fragment prepared from hybridoma cell line 11D3 (IgG2a, kappa) at 5 mg/mL mixed 1:1 with drop solution containing 30% PEG 400, 100 mM ADA pH 6.0 and 350 mM Li_2_SO_4_.

#### Data collection and structure determination

Before data collection, crystals were soaked in a cryoprotectant buffer containing 30% PEG 400 in the same crystallizing solution for 1 min, and rapidly frozen in liquid nitrogen. All diffraction data for drSLC38A9-Fab complex was collected at 100K using synchrotron radiation at the Advanced Photon Source (NE-CAT 24-ID-C and 24-ID-E). Diffraction data indexing, integration and scaling were performed with online server RAPD and CCP4 suite package *(35)*. Data collection statistics, phasing and refinement are given in Table S1. Molecular replacement using Phaser *(36)* was able to place two copies of Fab fragment (PDB ID: 1F8T) in native datasets. Helices of drSLC38A9 were manually placed in the density-modified map and extended within Coot *(37)* according to the reference model of ΔN-drSLC38A9-Fab complex (PDB ID: 6C08). Subsequent cycles of density modifications, model building and refinement were carried out in Phenix *(38, 39)* and Coot until structure completion (Fig. S2). The Ramachandran analyses of final structures were performed using Molprobity *(40)*. The model has been deposited into the PDB (PDB ID: 6dci).

#### Proteoliposomes reconstitution and radioligand uptake assays

The full-length drSLC38A9 and two variants, N-terminal deletion (truncate N-terminus from Met 1 to Val 96) and 5A (P79A, S80A, H81A, E82A, and Y85A) mutant protein, were expressed and purified as described above. Liposomes were prepared using a 3:1 ratio of *E. coli* total lipid extract (Avanti Polar Lipids) to chicken egg phosphatidylcholine (egg-PC, Avanti Polar Lipids) at 20 mg/mL in assay buffer (20mM MES pH 5.0, 150mM NaCl and 1mM DTT). An extruder with pore size of 0. 4 μm was used to obtain unilamellar vesicles. Triton X-100 was then added to the extruded liposomes at 10:1 (w:w) lipid:detergent ratio. Purified wild-type drSLC38A9 and variants were reconstituted at a 1:200 (w/w) ratio in destabilized liposomes and excess detergent was removed by SM2 Bio-Beads (Bio-Rad) at 4 °C overnight. Next day, proteoliposomes were collected, aliquoted and frozen at −80°C for storage until needed.

Transport reactions were initiated by adding [^3^H]-labeled amino acids (American Radiolabeled Chemicals) to 50 μL of 10-fold diluted proteoliposomes (total of 0.5 μg protein) to final concentration of 0.5 μM at room temperature. As controls, nonspecific uptake was assessed by using protein-free liposomes under identical conditions in parallel to experimental groups. At various time points, reactions were stopped by quenching the samples with 5 mL assay buffer followed by rapid filtration through 0.22μm membrane filter (GSWP02500, MilliporeSigma) to remove excess radioligands. The filter was then washed three times with 5 mL assay buffer, suspended in 10 mL of scintillation fluid and quantified by scintillation counting. A time course profile indicates that the retained radio-ligands reached saturation after 10 min. Measurements at various time points of the uptake were plot to establish the transport comparisons between various constructs of drSLC38A9. All experiment and control groups were repeated two to three times.

#### Co-Purification of zebrafish Rag GTPase complex with N-terminal fragment of drSLC38A9

The synthesized cDNA encoding RagA (UniProtKB - Q7ZUI2) and RagC (UniProtKB - F1Q665) from *Danio rerio* were cloned into pFastBac Dual vector. The Rag GTPase complex were overexpressed in *Spodoptera frugiperda* Sf-9 insect cells, which was harvested at 48 hours post-infection. Cell pellets were resuspended in lysis buffer containing 20 mM Tris pH 8.0, 150 mM NaCl. 30 homogenizing cycles were then carried out to break cells on ice, followed by a centrifugation at 130,000 *×g* for 30 mins. The supernatant was incubated with Ni-NTA Agarose (QIGEN) for 2 hours at 4°C. The resins were then washed with 5× column volumes of wash buffer containing 50mM Imidazole, 20 mM Tris pH 8.0, 150 mM NaCl. The protein was eluted by elution buffer containing 300 mM imidazole, 20 mM Tris pH 8.0, 150 mM NaCl, and then applied to gel filtration (Superdex 200 Increase 10/300 GL, GE Healthcare) in 20 mM Tris pH 8.0, 150 mM NaCl. The peak fractions were collected for further analysis.

To enhance solubility and stability, the N-terminal fragments of drSLC38A9 were fused with GB1 domain-tag *(41)*. drSLC38A9-N.1 is from Met 1 to Val 96, and drSLC38A9-N.2 is from Met 1 to Leu 70. The fusion proteins were overexpressed in *E. coli* BL21 (DE3) at 16°C for overnight with 0.2 mM isopropyl-β-D-thiogalactopyranoside (IPTG) as inducer. Then, the cells were harvested, homogenized in a lysis buffer containing 20mM Tris pH 8.0 and 150mM NaCl, and disrupted using a Microfluidizer (Microfluidics Corporation) with 3 passes at 15,000 p.s.i., followed by a centrifugation for 30 mins to remove cell debris. The supernatant was then loaded onto Ni-NTA Agarose and purified as above.

The purified Rag GTPase complex was mixed with excess GB1-drSLC38A9-N.x fragment at a molar ration of 1:2 for 1 hour, and the mixture was then subjected to gel filtration (Superdex 200 Increase 10/300 GL, GE Healthcare) in the buffer containing 20 mM Tris pH 8.0, 150 mM NaCl. SDS-PAGE and Coomassie blue staining was used to analyze the size exclusion chromatography elution profile.

All figures in this paper were prepared with PyMOL v1.8.6.0 *(42)*. Figure. S3 was prepared using the program Clustal Omega *(43)* for alignments and ESPript 3.0 *(44)* for styling.

**Table S1:** Data collection and refinement statistics

| S38A9-Fab<sup>†</sup> |  |
| --- | --- |
| **Data collection** |  |
| Space group | P 1 21 1 |
| Cell dimensions |  |
| *a*, *b*, *c* (Å) | 157.79, 82.51, 158.59 |
| α, β, γ (°) | 90.00, 106.02, 90.00 |
| Resolution (Å) | 49.37-3.40 (3.50-3.40)<sup>‡</sup> |
| R<sub>pim</sub><sup>¶</sup> | 0.084 (1.071) |
| Mean *I* / σ(*I*) | 6.5 (1.0) |
| Completeness (%) | 99.9 (99.8) |
| Redundancy | 11.8 (6.1) |
| CC<sub>1/2</sub> | 0.989 (0.350) |
| **Refinement** |  |
| Resolution (Å) | 49.37-3.40 (3.50-3.40) |
| No. of reflections | 54395 (5350) |
| R<sub>work</sub>/R<sub>free</sub> | 0.266/0.290<br>(0.352/0.368) |
| No. of non-hydrogen atoms |  |
| Protein | 12579 |
| Average *B*-factors |  |
| Protein | 106.96 |
| R.m.s. deviations |  |
| Bond lengths (Å) | 0.007 |
| Bond angles (°) | 1.33 |
| Ramachandran |  |
| favored (%) | 91.65 |
| outliers (%) | 1.38 |

<sup>†</sup> Four crystals were used to collect diffraction datasets which were processed, scaled, and merged using RAPD and AIMLESS.
<sup>‡</sup> Values in parentheses are for highest-resolution shell.
<sup>¶</sup> R<sub>pim</sub> is a measure of the quality of the data after averaging the multiple measurements and $R_{pim} = \frac{\sum_{hkl} [n/(n-1)]^{1/2} \sum_i |I_i(hkl) - \langle I(hkl) \rangle|}{\sum_{hkl} \sum_i I_i(hkl)}$ , where *n* is the multiplicity, *I<sub>i</sub>* is the intensity of the *i*<sup>th</sup> observation, <*I*> is the mean intensity of the reflection and the summations extend over all unique reflections (*hkl*) and all equivalents (*i*), respectively. (45)

**Fig. Sl.**
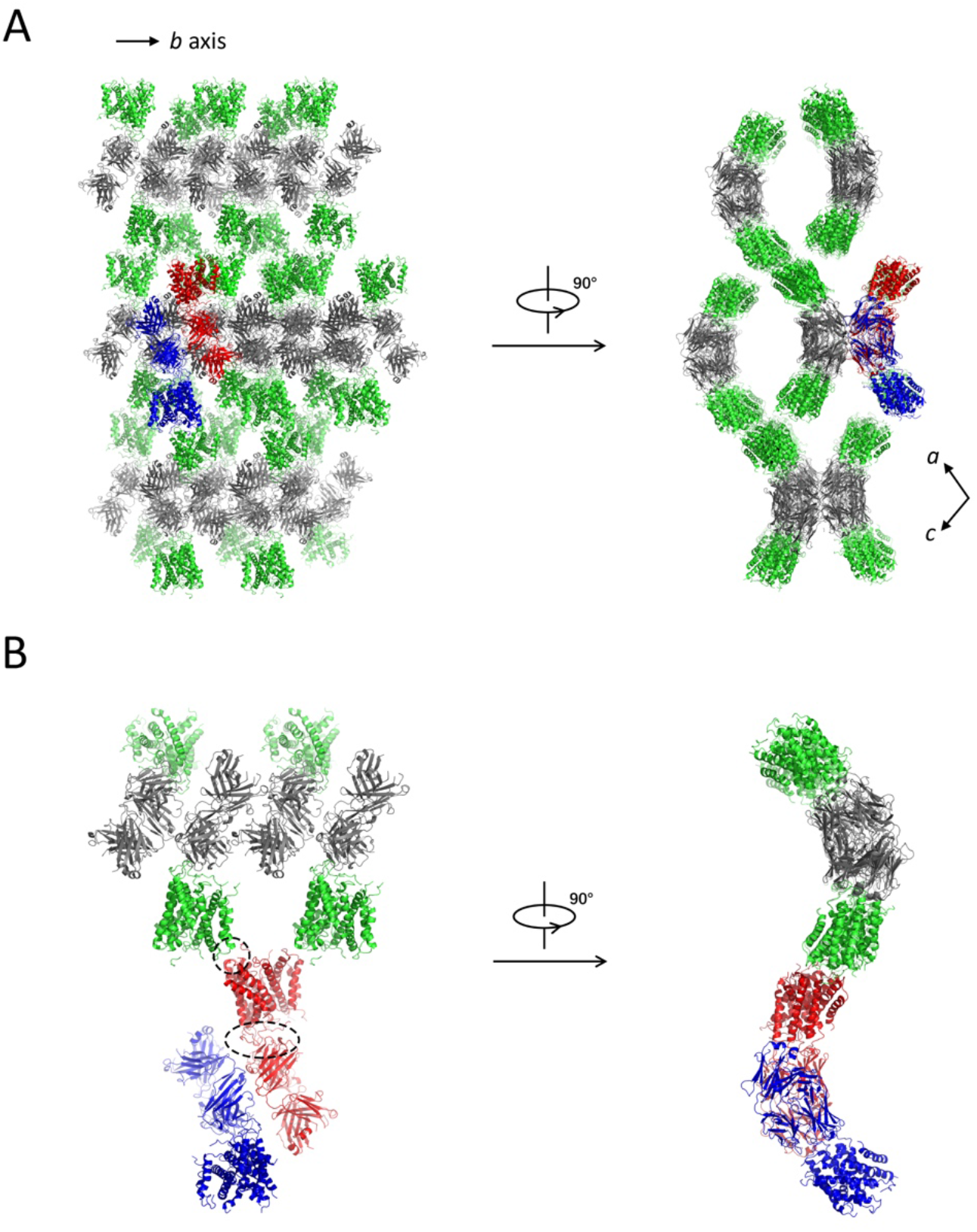
Crystal packing and asymmetric unit. (**A**) Crystal packing showing SLC38A9-Fab complex lattice. Fab fragments (grey) form continuous layers in the crystallographic *b* axis, which are connected by SLC38A9 (green) layers along the crystallographic *ac* plane in a propeller-like head-to-head manner. One asymmetric unit is selected to show the structural block comprising two-fold SLC38A-Fab molecules (red and blue). (**B**) Interactions between SLC38A9 molecules and Fab fragments. One SLC38A9 (red) makes biological contacts with the complementary determining regions (CDRs) of a Fab (red) by its luminal loops. It also has interactions between Loop 8-9 (red) and Loop 10-11 (green), which appear to be crystal contacts and non-specific.

**Fig. S2.**
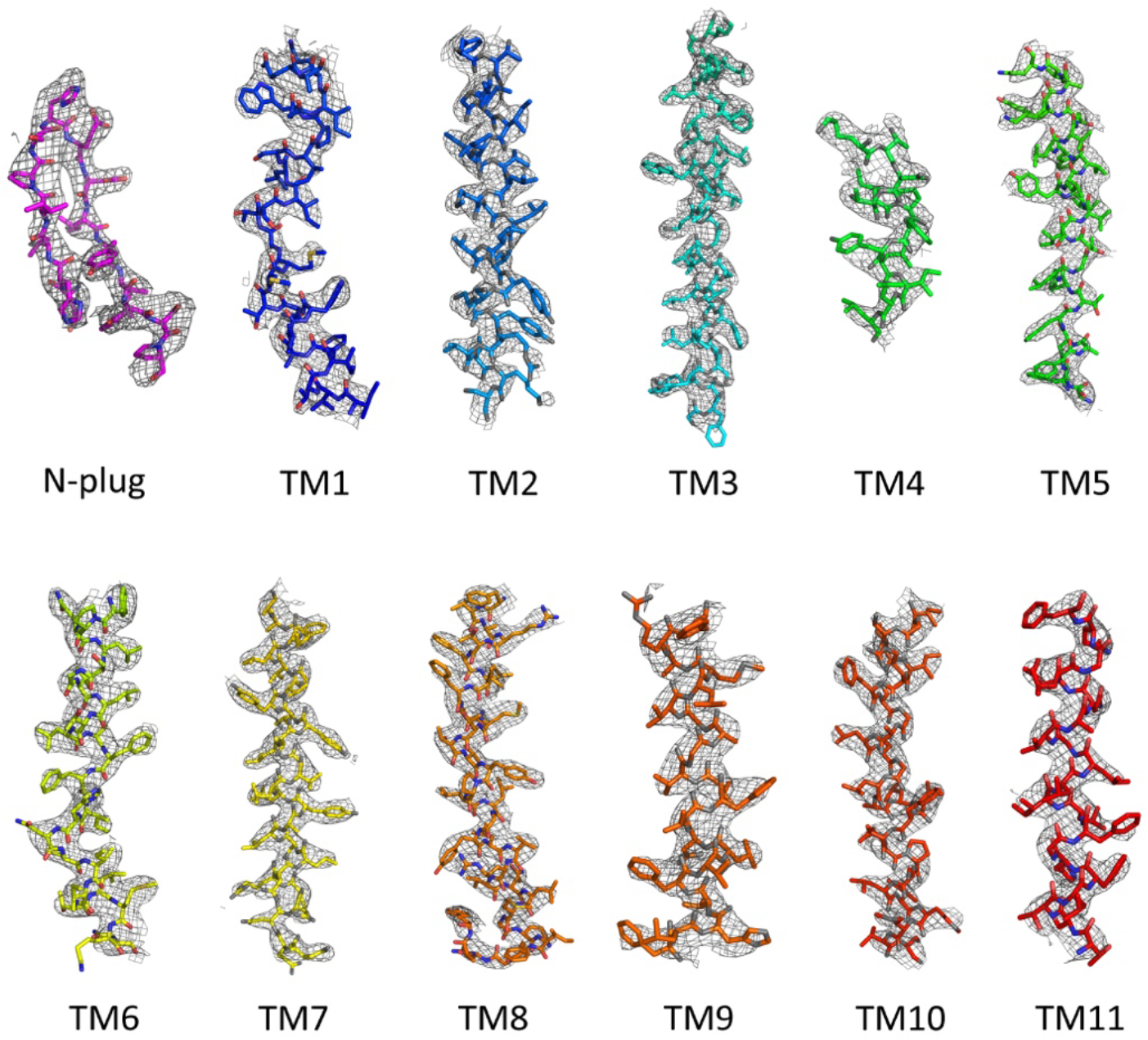
Overall experimental density of N-plug and membrane helices of drSCL38A9 are shown with 2Fo-Fc map contoured at 1.2 σ (gray mesh).

**Fig. S3.**
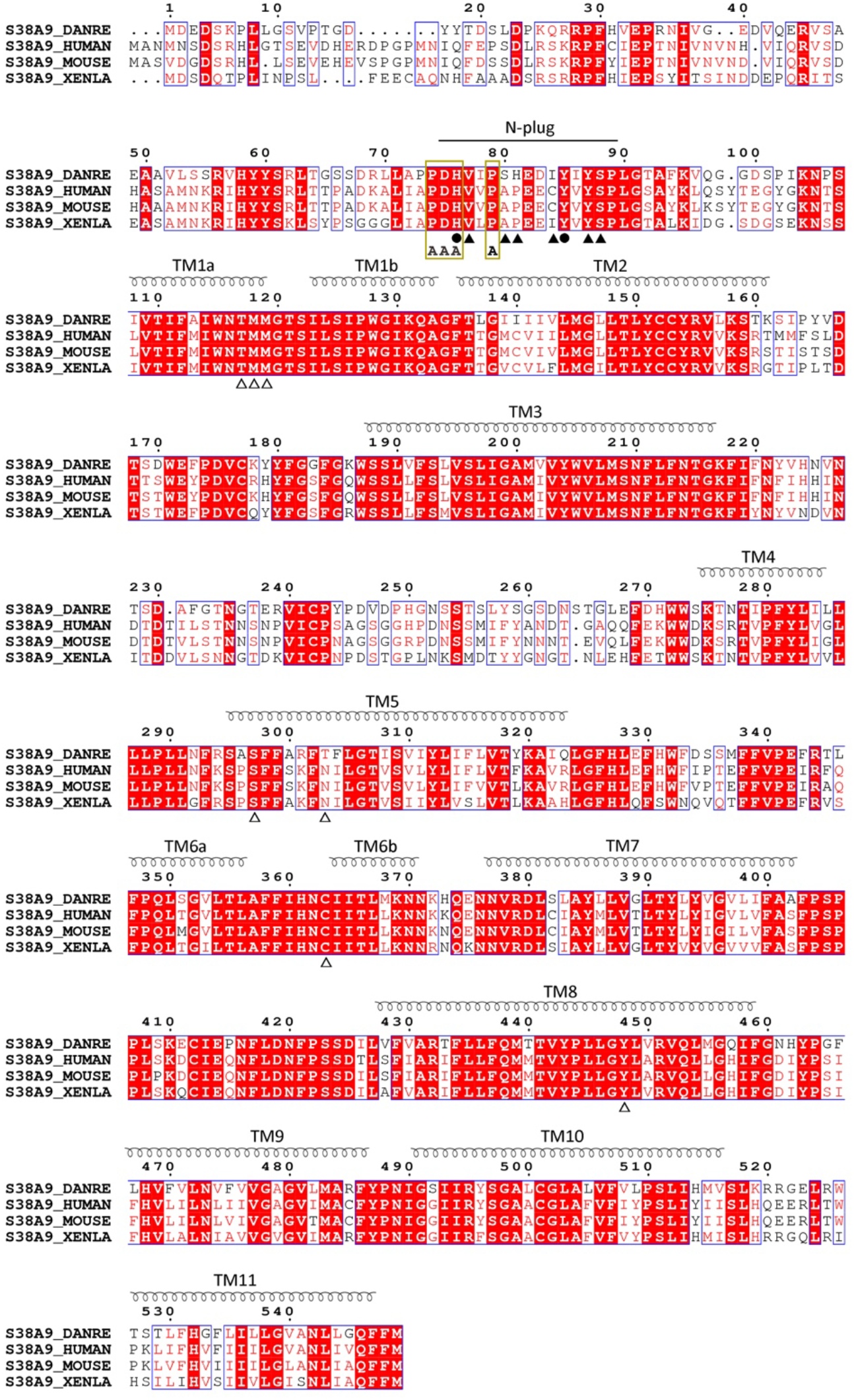
Sequence alignment of SLC38A9 from zebrafish, human, mouse and clawed frog. The N-plug of drSLC38A9 (from Asp 75 to Pro 89) is noted. Residues which have hydrogen bonds between the N-plug and transmembrane helices were labeled by triangles, while residues forming intra-interactions of N-plug were marked with circles. Alanine substitutions of residues in yellow box will abolish the binding of hSLC38A9 to downstream Rag GTPase complex.

**Fig. S4.**
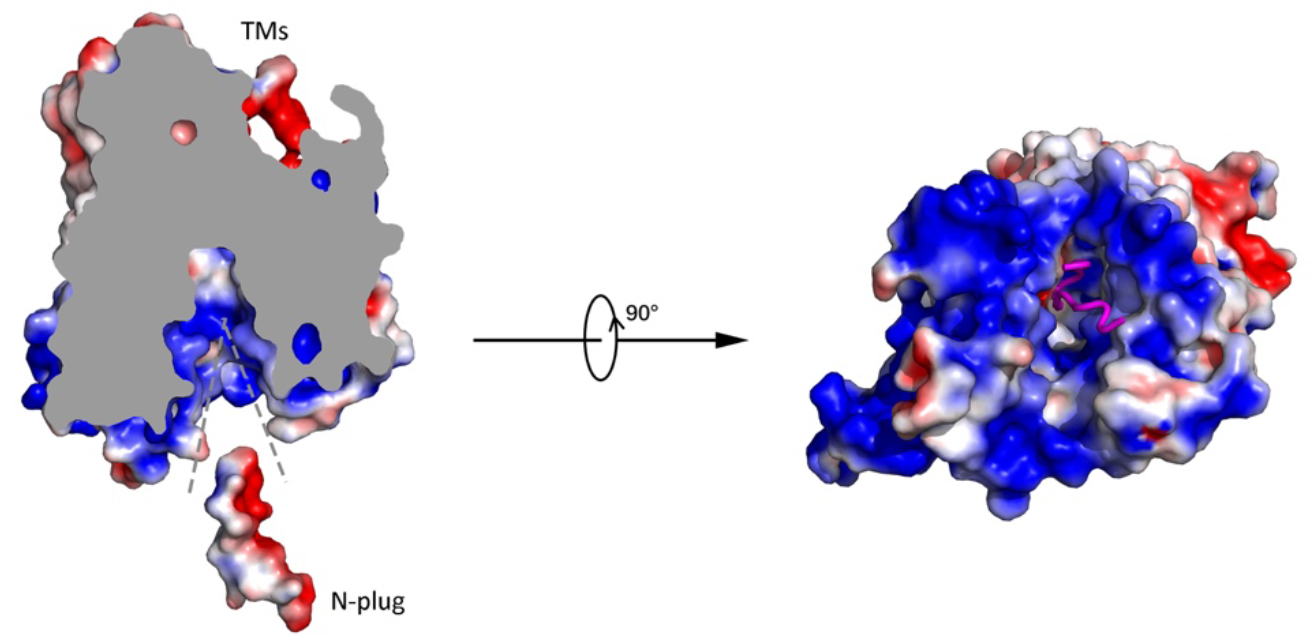
Electrostatic surface representation for drSLC38A9 showing the negatively charged N-plug blocks the access leading to cytosolic side, which is dominantly positively charged.

**Fig. S5.**
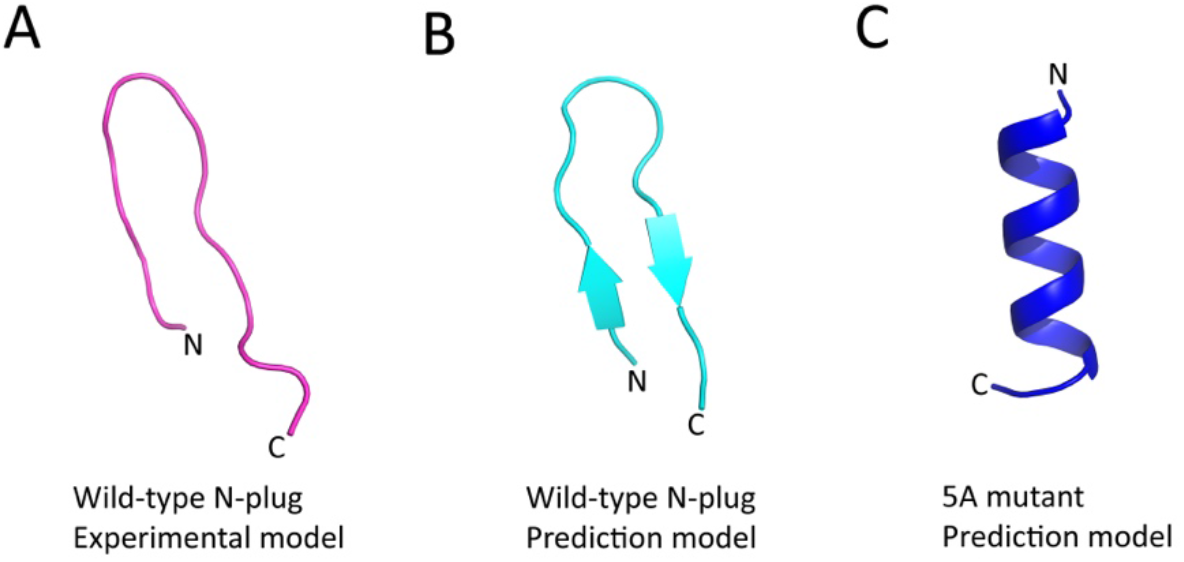
The 5A mutations (P79A, S80A, H81A, E82A, and Y85A) of N-plug disrupt its secondary structure. P79A, S80A, H81A, and E82A locate in the beta-turn motif. Y85A breaks hydrogen bonds of the beta-sheet network. (**A**) The experimental model of N-plug in current study. (**C** and **D**) The models of wild-type N-plug and 5A mutant peptide predicted by the same online sever PEP-FOLD 2.0. *(1)*

**Fig. S6.**
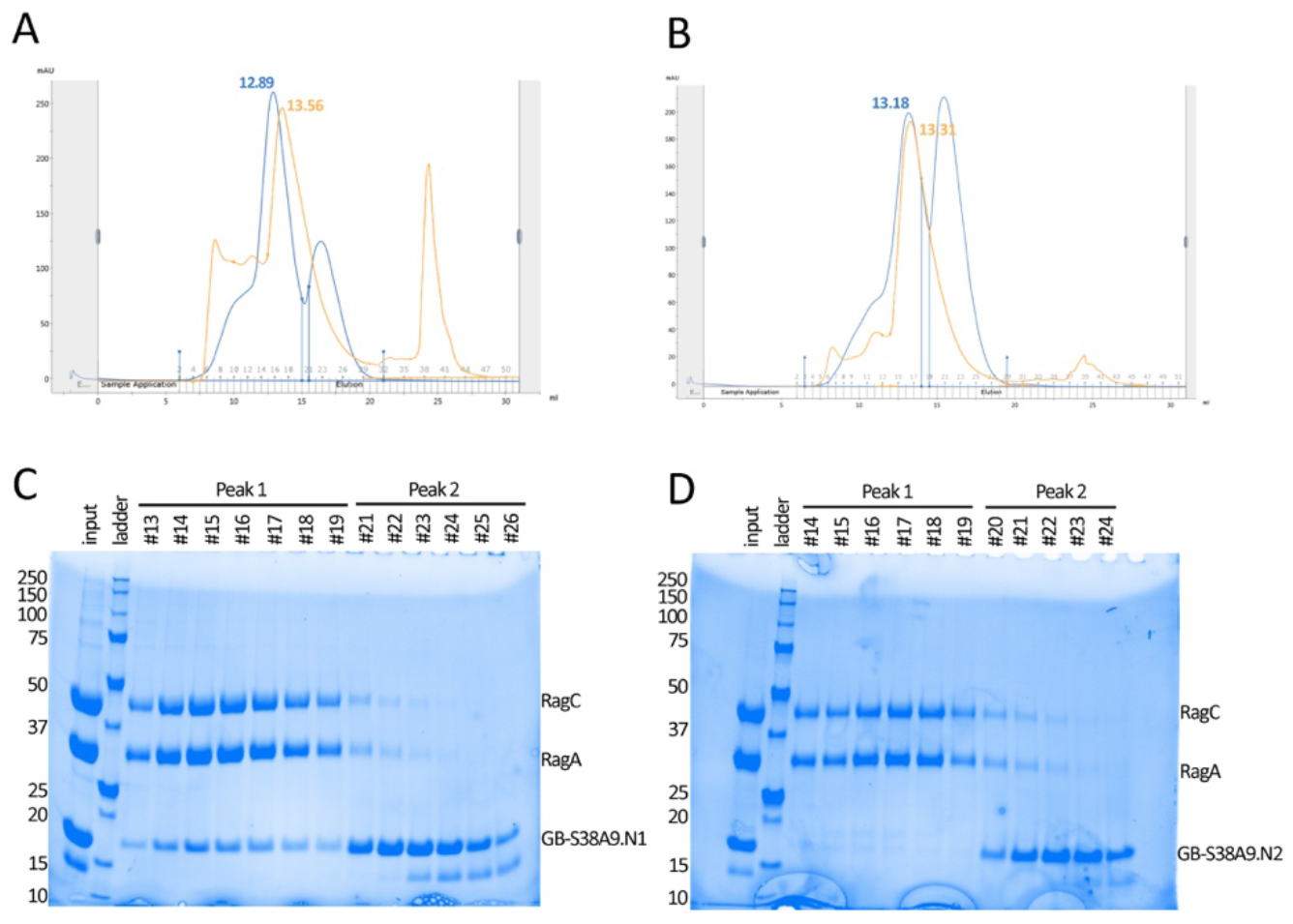
Co-purification of GB1 domain tagged N-terminal fragments of drSLC38A9 with zebrafish Rag GTPase complex. (**A**) The chromatogram displays a blue line with one peak at 12.89 mL retention volume (fractions 13-19) corresponding to drSLC38A9.N *(1–96)* and Rag GTPase complex and a second peak (fractions 21-26) corresponding to unbound N-terminal fragments. The orange superimposed curve depicts the zebrafish Rag GTPase complex eluted in the same column. Apparent peak shift was observed for the formation of drSLC38A9.N1 *(1–96)* and Rag GTPase complex. (**B**) Size exclusion chromatography profile of drSLC38A9.N1 *(1–70)* and Rag GTPase. No conspicuous peak shift was observed. (**C** and **D**) Fractions selected in (A) and (B) was sampled and analyzed on SDS-PAGE, including the input controls.

